# Decoding Age-specific Changes in Brain Functional Connectivity Using a Sliding-window Based Clustering Method

**DOI:** 10.1101/2022.09.27.509677

**Authors:** Aiying Zhang, David Pagliaccio, Rachel Marsh, Seonjoo Lee

## Abstract

Functional magnetic resonance imaging (fMRI) permits detailed study of human brain function. Understanding the age-specific development of neural circuits in the typically developing brain may help us generate new hypotheses for developmental psychopathologies. Functional connectivity (FC), defined as the statistical associations between two brain regions, has been widely used in estimating functional networks from fMRI data. Previous research has shown that the evolution of FC does not follow a linear trend, particularly from childhood to young adulthood. Thus, this work aims to detect the nuanced FC changes with age from the non-linear curves and identify age-period-specific FC development patterns. We proposed a sliding-window based clustering approach to identify refined age interval of FC development. We used resting-state fMRI (rs-fMRI) data from the human connectome project-development (HCP-D), which recruited children, adolescents, and young adults aged from 5 to 21 years. Our analyses revealed different developmental patterns of resting-state FC by sex. In general, females matured earlier than males, but males had a faster development rate during age 100 -120 months. We identified four developmental phases: network construction in late childhood, segregation and integration construction in adolescence, network pruning in young adulthood, and a unique phase in males -- U-shape development. In addition, we investigated the sex effect on the slopes of FC-age correlation. Males had higher slopes during late childhood and young adulthood. These results inform trajectories of normal FC development, information that can in the future be used to pinpoint when development might go awry in neurodevelopmental disorders.

**Highlight:**

- Propose a novel sliding-window-based framework to identify refined age intervals of functional connectivity (FC) development.
- Identify four developmental phases: network construction in late childhood, segregation and integration in adolescence, network pruning in young adulthood, and a unique phase in males -- U-shape development.
- Characterize the representative FC pattern for each developmental phase based on global network statistics, modular connectivity, and hub ROIs.
- Reveal sex differences in developmental timing, rate, and patterns of resting-state FC.

## 1. Introduction

Advances in neuroimaging methodologies during the past two decades permit detailed study of human brain maturation. Investigators have increasingly focused on understanding dynamic changes in brain structure and function over development, particularly as many psychiatric disorders originate in childhood [1]. Therefore, establishing typical patterns of age-related changes in functional neural circuits could be a significant step in the translational application of neuroimaging. Such established patterns could be used as reference data to identify when developmental abnormalities arise and, further, to determine where, when, and how to intervene to prevent illness persistence [2]. These patterns are also crucial for generating hypotheses regarding the neural bases of developmental psychopathologies [3].

Maturation of functional connectivity (FC) can be examined by quantifying age-related changes in the strength and spatial distribution of intrinsic brain networks [4, 5]. Resting-state functional MRI (rs-fMRI) measures the intrinsic activity of the brain [6] and potentially provides a broader window into core features of the brain’s functional organization [7]. A recent review [8] of FC changes at rest over the human lifespan suggested that the development of FC does not follow a linear trend, particularly during adolescence, and the onset and the duration of the changes may vary from region to region. Therefore, previous methods using predefined age-related groups [9, 10, 11] or linear regression [11, 12] may not be sensitive to capture non-linear FC development.

One potential solution is to use non-linear curve fitting methods to probe associations between FC and age. Various methods have been proposed in the literature, such as polynomial or spline regression, and have been applied in brain trajectory studies [8, 13]. However, it can be difficult to interpret FC-age associations as these curves become increasingly complicated. Consequently, these methods may not sensitively capture the FC change structure in terms of onset, duration, and magnitude without additional estimations. To address this limitation, we directly estimate the slope of curve instead of delineating the curve itself, which has a clearer interpretation. For example, when age has no association with FC, then the slope equals zero. Alternatively, when changes in age associate with a large change in FC, then the absolute value of the slope is large.

On the other hand, sex differences have been well-recognized as significant contributor in the cognition development [14] and psychiatric disorders [15]. Previous neuroimaging studies have examined FC differences by sex and linked them with various cognitive domains, such as language [16], motor skills and spatial ability [14]. Further, neurodevelopmental disorders, including intellectual disability (ID), autism spectrum disorder (ASD), and attention-deficit hyperactivity disorder (ADHD), are more prevalent in males [17, 18]; other disorders such as depression and anxiety disorders are more prevalent in females [19]. In addition, some studies suggest different onsets of psychiatric illnesses in boys and girls, for example, conduct disorder [20]. Studies have suggested that there is robust difference in FC distinctiveness trajectories in youth and delayed stabilization in FC are related to psychiatric disorders [21]. However, it remains unknown whether there are robust sex differences in terms of FC development curve.

This study proposes a data-driven approach to reveal functional networks that share similar FC-age association patterns over development. Specifically, we propose a sliding-window-based clustering approach to identify refined age intervals of change in FC. We used rs-fMRI data from the Human Connectome Project-Development (HCP-D) study, which aims to recruit a total of 1300 children, adolescents, and young adults ranging in age from 5 to 21 years [22]. We analyzed FC-age associations separately by sex and then characterized each identified FC pattern based on three aspects: global network statistics, modular connectivity, and hub regions-of-interest (ROIs). Global network statistics describe important properties of brain networks with a small number of neurobiologically meaningful and easily computable topological measures. Modular connectivity reflects the intra- and inter-network associations. Hub ROIs are the most important centers in a FC network.

## 2. Result

As illustrated in Figure 1, we used the proposed sliding-window-based clustering approach to characterize FC development in 528 youth ages 5 to 21 years imaged with rs-fMRI on two days as part of the HCP-D study. We present findings consistent across day 1 vs day 2 scans.

**Figure 1:**
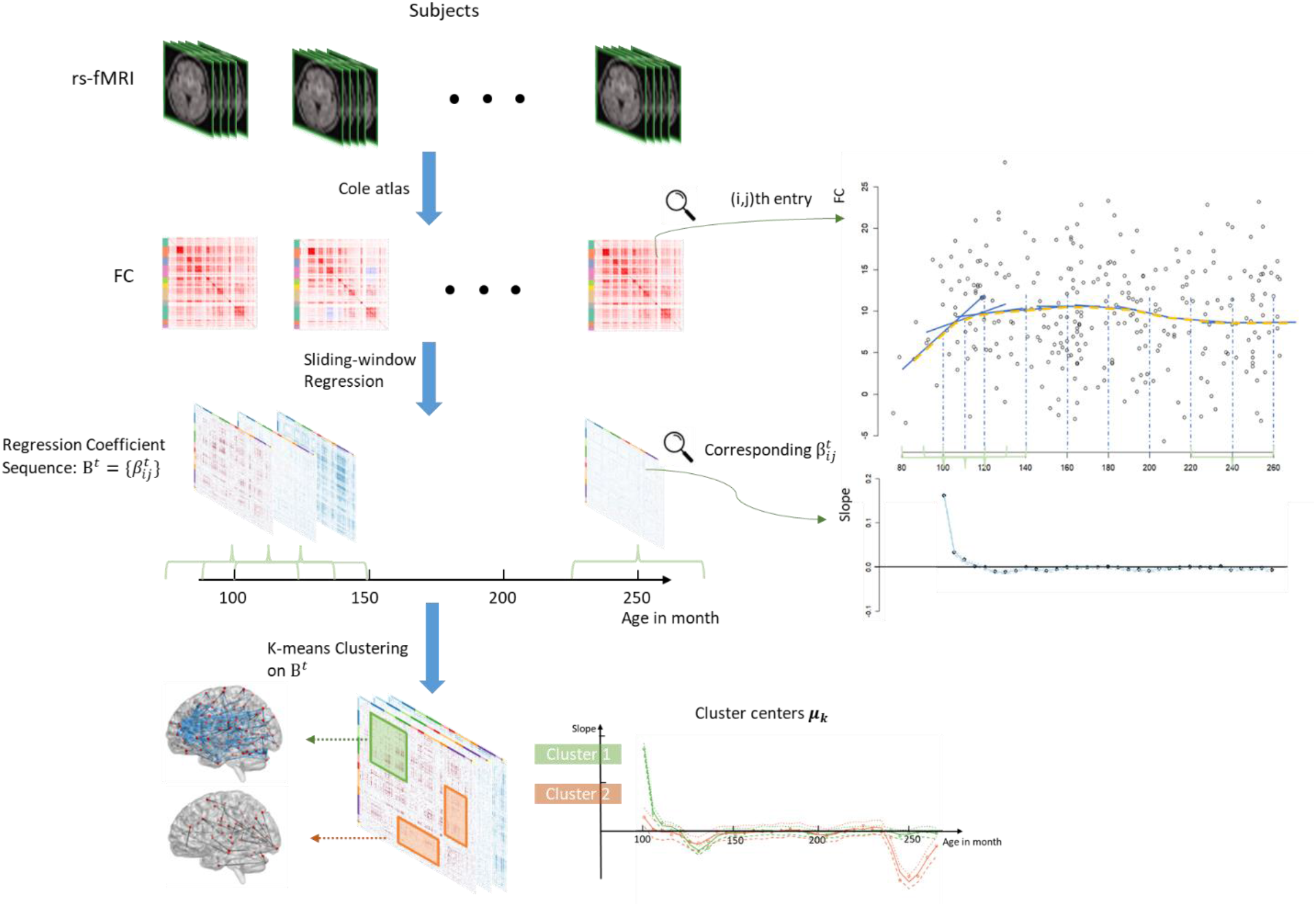
The flowchart of detecting age-specific development patterns.

### 2.1 Association of age with FC by sex to identify critical development periods

Figures 2-3 show the identified cluster centers of the regression coefficient sequences (in solid lines) and representative shapes (in dashed lines) of the age-FC associations. For example, Cluster F1 in Figure 1 shows positive regression coefficients early in development (solid line) reflecting a positive slope between age and FC; this is shown in the dashed line where connectivity values increase across early development. We tested on the validation dataset and found consistency of the cluster curves (in Figure S3).

**Figure 2:**
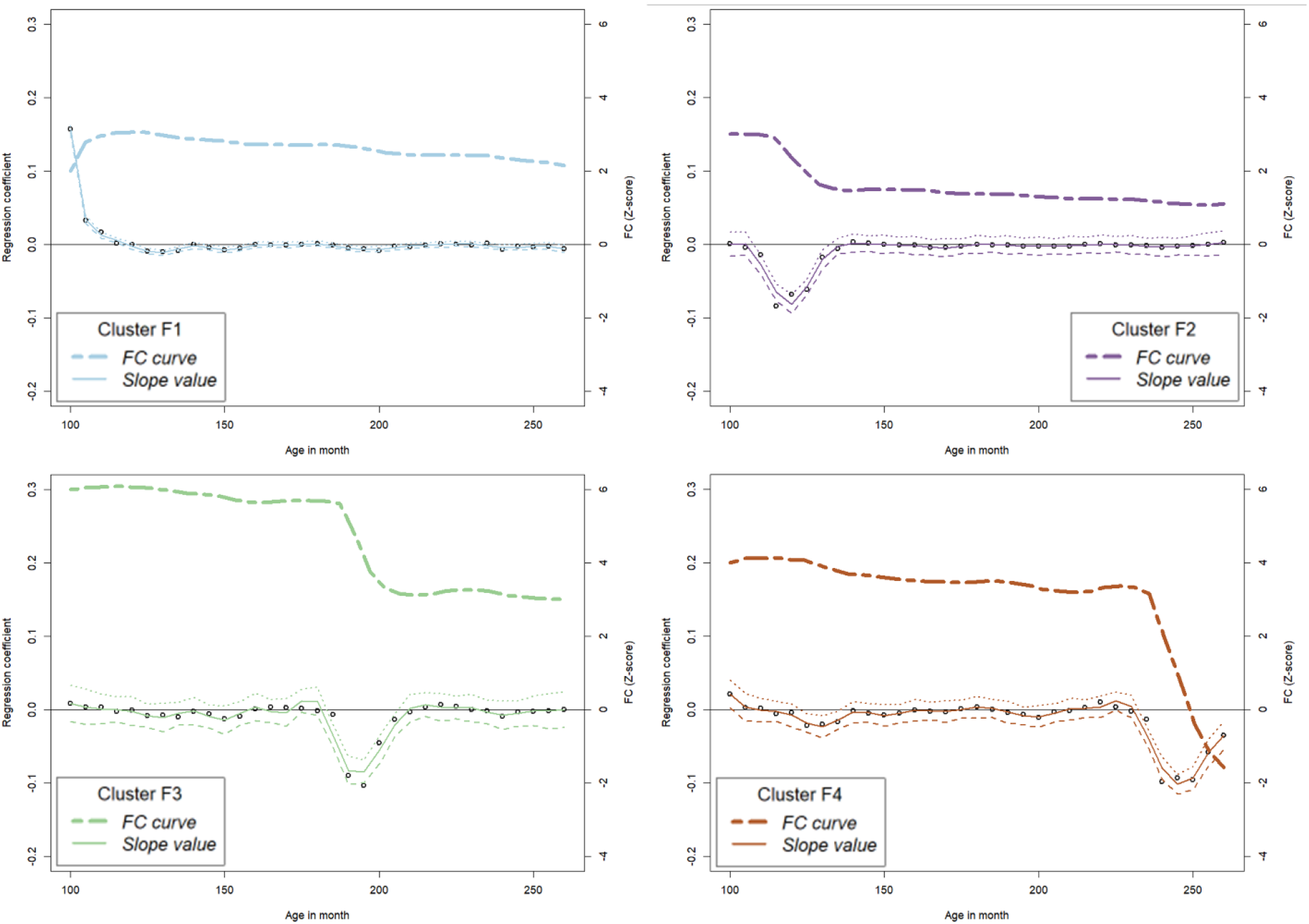
The developmental curves clustered by K-means for females. The solid line represents the sliding-window regression coefficient sequence with 95% confidence interval, and the dashed line represents the shape of the FC-age association in each cluster.

**Figure 3:**
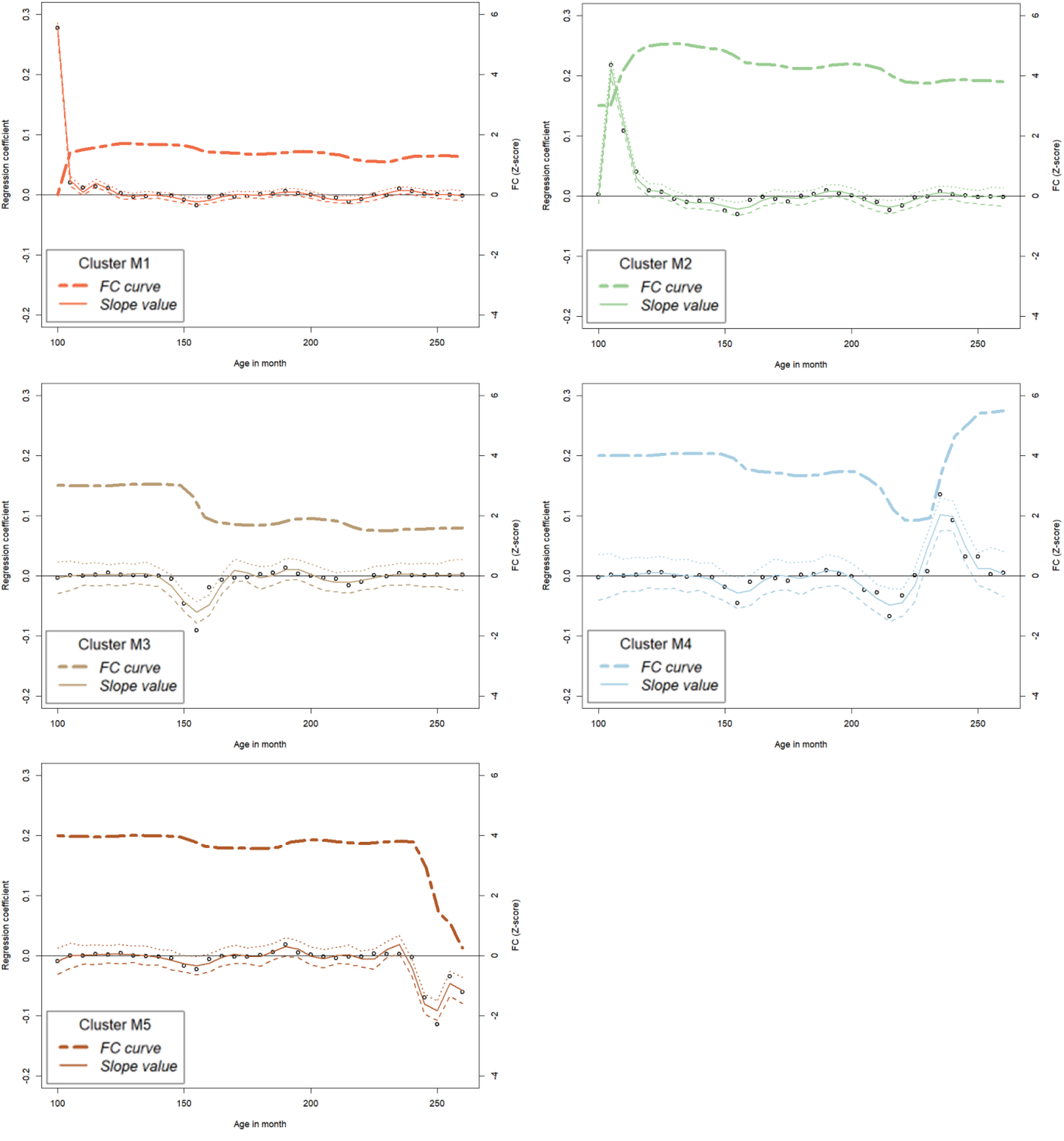
The developmental curves clustered by K-means for males. The solid line represents the sliding-window regression coefficient sequence with 95% confidence interval, and the dashed line represents the shape of the FC-age association in each cluster.

#### Similarities

Males and females both exhibited one cluster positive FC-age associations in late childhood (F1 in Figure 2 and M1 in Figure 3) and one with negative FC-age associations in young adulthood (F4 and M5).

#### Differences

Females started to show negative FC-age associations in early adolescence (F2) and had negative FC-associations afterwards (F2 - F4). Males showed negative FC-age associations (M3) later than females and had positive FC-association before (M2) and after (M4).

### 2.2 FC patterns of development as potential development markers of each age period

For each cluster of FC and age association curves, we extracted their FC patterns. Detailed information was provided in Figures S5-S15. We showed the network statistics of the FC pattern from each representative curve in Table 1 and summarized the corresponding developmental characteristic by sex. The cluster names in Figures 2&3 and Table 1 are matched, as well as in the rest of the paper. Based on the period of change and the developmental characteristic, we identified four phases of development.

**Table 1:** Summary of the representative shapes of the association of age with FC.

| **Female** |  |  |  |  |  |  |
| --- | --- | --- | --- | --- | --- | --- |
|  | Age period (months) | Max slope | Density | Transitivity | Global Efficiency | Developmental Characteristic |
| F1 | [100,110] | 0.157 | 0.0030 | 0.0785 | 0.0600 | Segregation |
| F2 | [110,130] | -0.084 | 0.0084 | 0.0336 | 0.1955 | Integration |
| F3 | [190,205] | -0.103 | 0.0090 | 0.0857 | 0.1279 | Integration and segregation |
| F4 | [235,260] | -0.098 | 0.0008 | 0.0128 | 0.0107 | Segregation |
| **Male** |  |  |  |  |  |  |
|  | Age period (months) | Max slope | Density | Transitivity | Global Efficiency | Developmental Characteristic |
| M1 | [100,110] | 0.277 | 0.0001 | 0 | 0.0002 | Connectivity construction |
| M2 | [105,120] | 0.218 | 0.0003 | 0.0414 | 0.0011 | Segregation |
| M3 | [150,160] | -0.091 | 0.0079 | 0.0433 | 0.1576 | Integration and segregation |
| M4 | [210,250] | - | 0.0011 | 0.0118 | 0.0133 | U-shape development in segregation |
| M5 | [240,260] | -0.115 | 0.0005 | 0 | 0.0062 | Integration |

#### Network construction in late childhood – Cluster F1, M1, M2

Two curves in males were detected in this phase. To separate them, we called M1 first construction and M2 second construction. The FC patterns in both sexes involved the visual network (VN: V1N and V2N) and default mode network (DMN): for females, the functional connections appeared on both cortical and subcortical areas in F1; for males, they mainly appeared on subcortical areas in the first construction and on cortical areas in the second construction.

For females, the hubs were mainly located in the thalamus and the frontal and temporal lobes. For males, the hubs were mainly located in the putamen and hippocampus (first construction), the frontal gyrus, hippocampus and temporal gyrus (second construction).

#### Segregation and Integration in adolescence – Cluster F2, F3, M3

Two curves were detected in females during adolescence: F2 in early adolescence ([110, 130]) and F3 in late adolescence ([190, 205]). The FC patterns in both sexes showed segregation and integration construction in VN and cingulo-opercular network (CON). In early adolescence (F2), the FC patterns mainly appeared in the subcortical VN, CON, frontoparietal network (FPN) and ventral-multimodal network (VMN). In late adolescence (F3), the FC patterns mainly appeared in the cortical CON, cortical auditory network (AUD), and subcortical VN. For males, the FC patterns mainly appeared in the VN, CON, and cortical DMN.

For females, the hubs were located in the temporal lobe, basal ganglia and thalamus in early adolescence; in the right thalamus, cerebellum, and temporal lobe in late adolescence. For males, the hubs appeared to be mainly in the caudate, thalamus, left cerebellum, and right temporal lobe during both adolescent periods.

#### U-shape development in male – Cluster M4

This was a unique development phase for males which appeared as a strengthened U-shaped association between FC and age. The FC patterns mainly involved the intra- and inter-connectivity in AUD. The corresponding hubs appeared at the thalamus and the vermis 6 area in the cerebellum.

#### Network pruning in young adulthood – Cluster F4, M5

The FC patterns in both sexes mainly appeared in subcortical areas: for female, it mainly involved the intra-modular connectivity in V1N and V2N; for males, it mainly involved the FPN and language network (LAN). The hubs for both FC patterns were mainly located in the cerebellar areas.

### 2.3 Age-related changes in modular connectivity

As shown in Figure 4, sex differences were noted at the modular level. In general, females had earlier modular connectivity development than males. The age interval for females was [100, 205] and for males was [150, 260]. However, males showed an earlier development phase than females involving CON, DMN and V2N.

**Figure 4:**
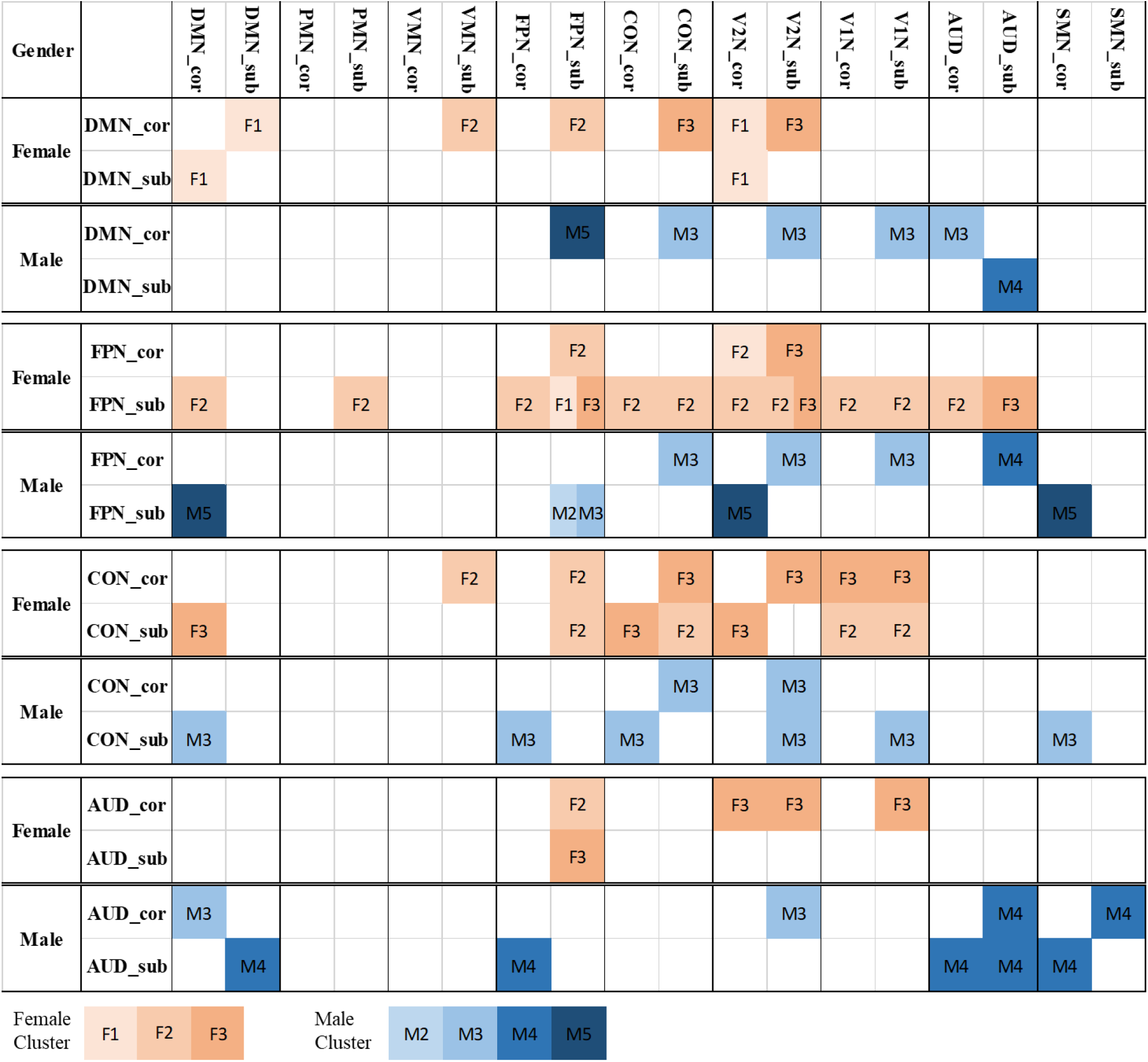
The significant intra- and inter- modular connectivity of selected functional network by sex (female: in orange, male: in blue). The lightness of the color represents the age period of the cluster. The significant modular connectivity of all functional networks can be found in the Supplementary file S14, S15.

#### CON

The modular connectivity change in CON appeared in the adolescent phase for both females (F2, F3) and males (M3). Three pairs of inter-modular connectivity: cortical-CON ↔ subcortical-CON, cortical-CON ↔ subcortical-V2N, subcortical-CON ↔ cortical-DMN, had significant changes in both sexes, which appeared at [150,160] in males and [190,205] in females.

#### FPN

Females had active and consistent inter-modular changes in the subcortical-FPN, mostly during early-adolescence, while males showed an association with age at a later period ([150,260]). The intra-connectivity of subcortical-FPN showed two significant phases of change in both sexes with females showing early changes in childhood and males showing early changes in adolescence.

#### AUD

Females showed a significant decrease in interconnectivity in late adolescence (F3) while males had a unique U-shaped development phase in late adolescence to young adulthood.

#### DMN

In females, childhood network construction (F1) appeared in the interconnectivity in DMN ↔ cortical-V2N and cortical-subcortical DMN. For the interconnectivity in cortical-DMN ↔ subcortical-V2N, and cortical-DMN ↔ subcortical-CON, males showed an earlier development phase than females.

### 2.4 Sex effect on the slopes of FC development

We observed 2 significant sex differences in the slopes of age-related FC where males had higher slope: one during late childhood (age interval [115,130], max regression coefficient 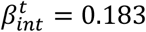) and the other during young adulthood (age interval [230,260], max regression coefficient 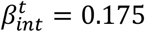). The FC patterns of each age period with pronounced sex differences in slope can be found in the Supplementary file (Figures S16, S17).

Figure 5 visualizes the hub ROIs in each age period with slope differences. For the late childhood period, the identified hubs, mainly located in the left temporal lobe and caudate, functionally belong to the CON, DAN and AUD. For the young adulthood period, the identified hubs, mainly located in the right parahippocampus, right postcentral gyrus and the cerebellum, functionally belong to the SMN and V2N. Detailed information about the hubs is provided in the Supplementary file (Table S1).

**Figure 5:**
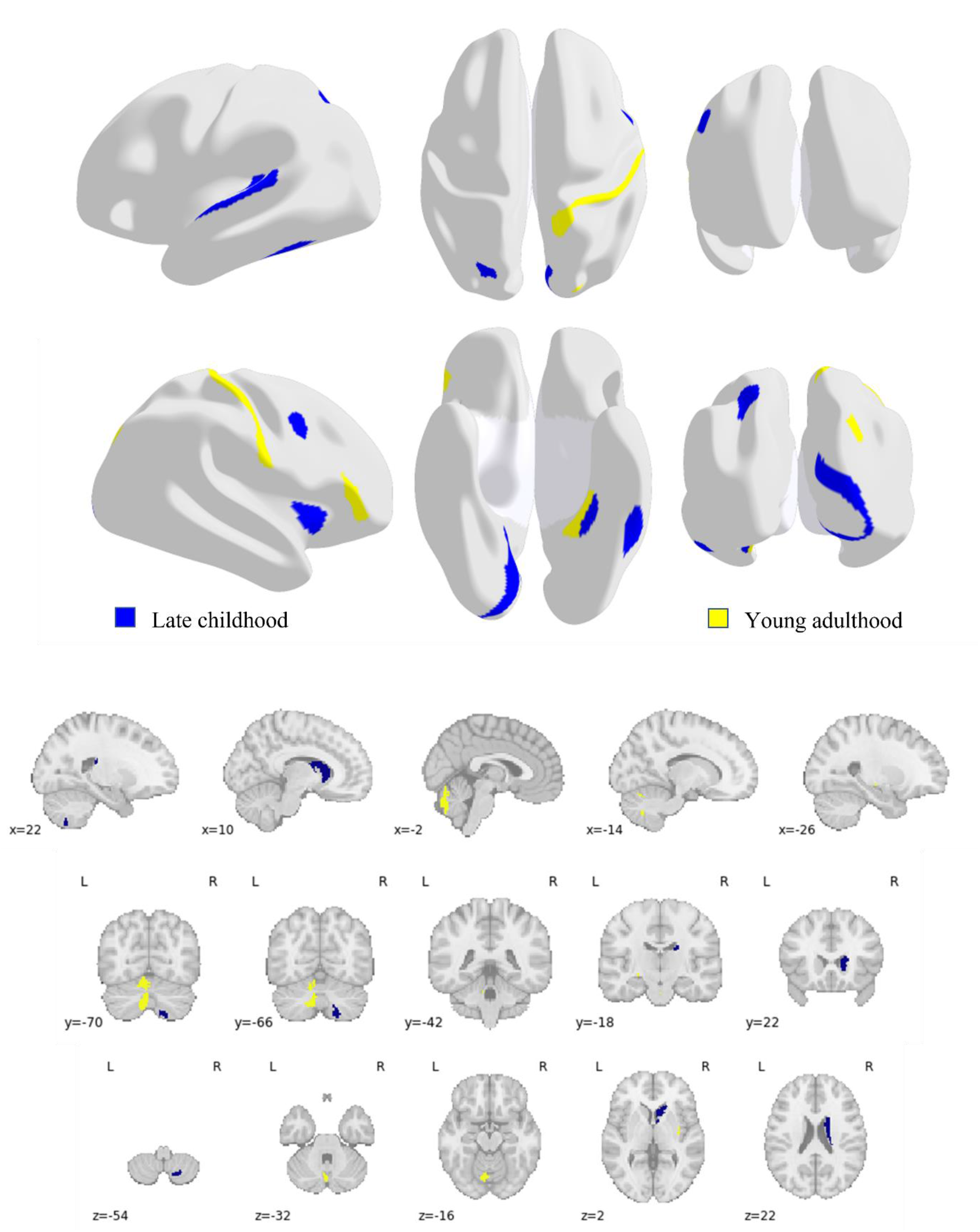
Visualizations of the hub ROIs that have significant slope differences in sex on the cortical surface and subcortical areas.

### 2.5 Sensitivity Analysis

We further conducted sensitivity analysis using various measurements of the FC-age association, namely, Pearson correlation, Spearman correlation, linear regression with no covariates, linear regression with motion correction. The results in Figure 6 indicate that the identified developmental curves are stable over different measures.

**Figure 6:**
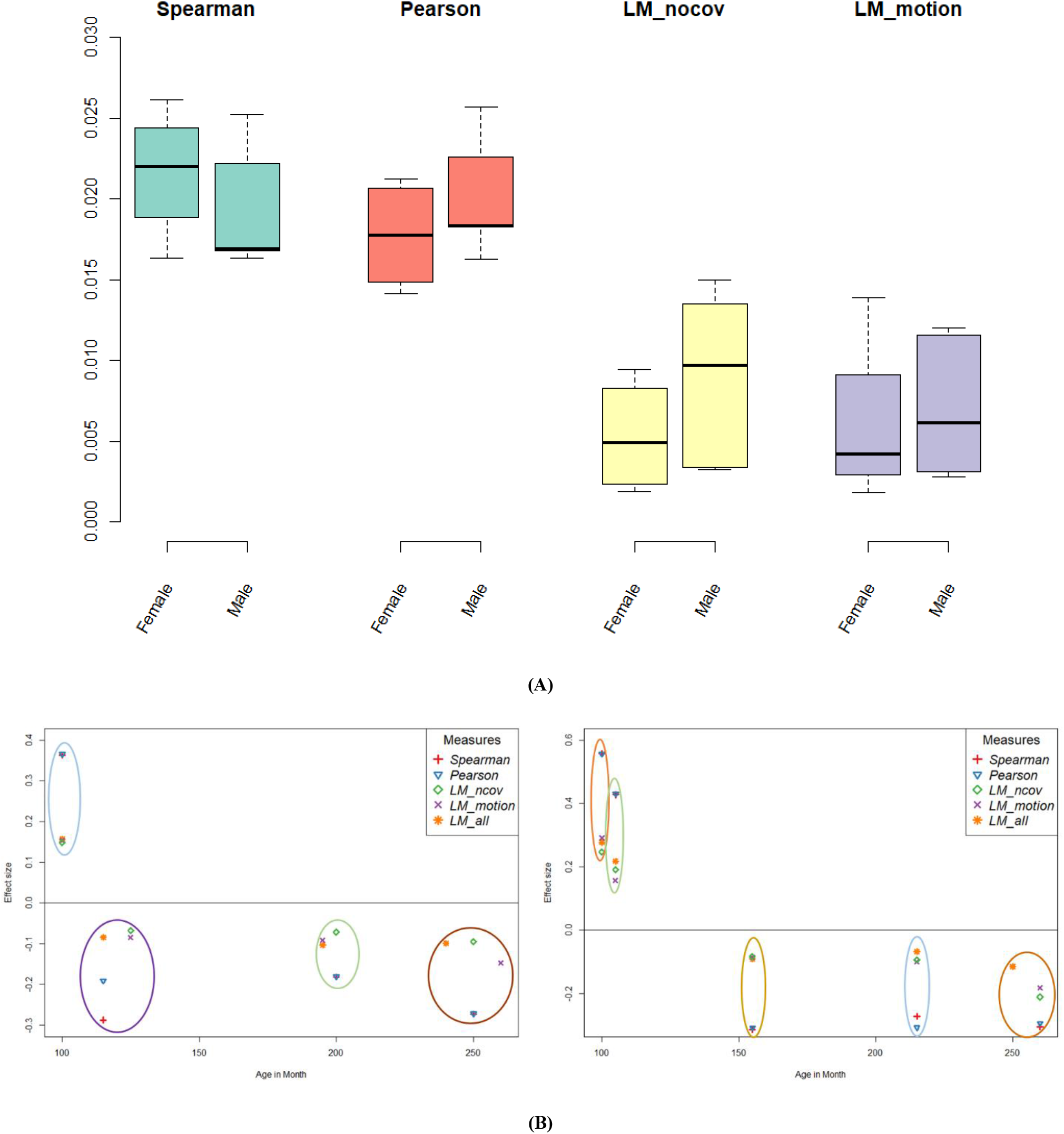
Sensitivity analysis using various measures. (A) Boxplots of the Euclidian distances of the coefficient curves of selected measures, i.e., Spearman coefficient, Pearson coefficient, linear regression with no covariate (LM_ncov), linear regression with motion correction (LM_motion), versus linear regression with covariates (LM_all). (B) The cluster centers under various measures. The max slope values were visualized.

## 3. Conclusion and Discussion

Our analyses reveal developmental patterns of resting-state FC by sex. We showed sex differences in FC-age associations such that females matured earlier than males, but males had faster rate of development. These findings are consistent with those of many studies showing robust sex differences in functional brain organization [21, 23, 24]. We identified four phases of development, which are network construction in late childhood, segregation and integration in adolescence, network pruning in young adulthood, and a unique phase of U-shape development in males.

Network construction mainly involved the visual and default mode networks. Multiple studies have identified visual related development across childhood and adolescence using tasks of visual working memory [25, 26], visual word processing [27], visual face recognition [28], etc. For instance, improved performance on a standard visual working memory (VWM) test [29, 30] has been shown from late childhood to early adolescence and the VWM continues to develop through adolescence [25]. In this study, we contribute to understanding the mechanistic underpinnings of the visual-related development. We also found significant modular connectivity within DMN in females of this phase. Previous studies show that the connectivity strength with the DMN is greater in adolescence than in children and is greater in adults than in children [31]. Our findings suggested that the connectivity strength difference between childhood and later period may be ascribe to the late-childhood construction.

The segregation and integration phase in adolescence is the period with major changes occurring across the brain at many different levels of brain functioning. Significant changes in modular connectivity in CON appeared in this phase for both females and males. Females had active and consistent inter-modular changes in the subcortical-FPN during early-adolescence development phase. Studies reported sex differences in neural activation during cognitive control tasks in FPN, in which adolescent and adult males display stronger activation than females [32, 33]. Based on our analyses results, this may be ascribed to the earlier adolescent phase adolescence construction in females.

U-shape development in males mainly involved the intra- and inter- connectivity in AUD, while the AUD development in females appeared at the late-adolescent phase. Studies [34, 35] demonstrated that cortical auditory circuits continue to mature during adolescence. We further provided the detailed onset of AUD development and age-related changes in these circuits for each sex.

In general, modular connectivity developed earlier in females than males. For example, significant modular connectivity between cortical-DMN and subcortical-FPN was detected in early adolescence for females and in young adulthood for males. However, males showed earlier development of modular connectivity with CON and DMN than females. The CON, FPN and DMN support higher-order cognitive abilities and alterations in these networks are associated with both cognitive impairments [36] and neurodevelopmental diseases that are characterized by such impairments [37, 38, 39]. In addition, cognitive control processes including the CON and FPN have been found to undergo substantial maturation during this period, with protracted rates of development of parietal and prefrontal cortices [40] likely reflecting synaptic pruning and neuronal specialization [41]. Studies have shown that ineffective reduction of functional connectivity between the DMN and FPN during cognitive control can interfere with performance in healthy individuals — a phenomenon present in psychiatric disorders, such as depression [42]. The disruption of inter-connectivity disproportionately affects girls and young women [43], consistent with the higher prevalence of depression in females. Our results indicated that functional changes across DMN appeared much earlier in females than males, which provided a possible explanation for heightened vulnerability for depression in adolescent girls.

We also investigated sex effects on FC development. Two significant differences in the slopes of age-related FC across sexes emerged with males having a higher slope during late childhood, corresponding to difference of segregation development in male and integration development in female; the other one was during young adulthood, corresponding to the U-shape development in male. For the late childhood difference, the identified hubs, mainly located at the left hemisphere on temporal lobe; while for the young adulthood difference, hubs were at the right hemisphere on postcentral lobe.

In addition, the cerebellum emerged as a hub area in various phases for both sexes after early adolescence (female: F3, F4; male: M3, M4, M5). The cerebellum is recognized as a prominent role in higher cognitive functions involving language, emotion regulation, learning and working memory [44, 45]. Evidence shows that during adolescence, subdivisions of the cerebellum have distinct developmental trajectories in late puberty [46], which aligns with our results.

Our findings provide information on the mechanisms that underlie the FC-age associations in healthy populations. The value of our work is that we established a reference for age-related FC development from childhood to adulthood, where deviations from an expected range can be used to trigger further investigations or interventions. Direct comparison with normal FC development may help 1) understanding of some neurodevelopmental disorders such as ADHD, OCD, or autism [3], 2) identify subtle changes early in the disease course, 3) characterizing patient– specific abnormalities.

## 4. Materials and Methods

### 4.1. Participants

We used resting-state fMRI (rs-fMRI) data from the HCP-D study, which extended the adult HCP study to examine the brain connectome in healthy children acquired at four sites across the USA [22]. A total of 1300 children, adolescents, and young adults ranging in age from 5 to 21 years will be enrolled with the following exclusion criteria of individuals who a) could not complete the study, b) had health problems that would compromise their inclusion or jeopardize their anonymity, c) neurodevelopmental disorders, and/or d) contraindications to MRI. In the current release (version 2.0), the baseline data from N=625 participants were available. We excluded participants with excessive head motion during scanning (see further). The final sample included 528 participants.

### 4.2. Image acquisition and pre-processing

During HCP-D rs-fMRI scanning, participants were instructed to stay still, stay awake, and blink normally while looking at the fixation crosshair. For participants 8 years and older, 26 min of resting state scanning were acquired in four runs over two consecutive days, using the 3T Siemens Prisma platform with the following parameters TR/TE= 800/37 ms, flip = 52, FOV= 208 × 180 mm, matrix = 104 × 90, slices = 72, voxel size = 2.0 × 2.0 × 2.0 mm. For the youngest ages (5–7 years), the individual runs were reduced to 3.5 min each because young children generally cannot tolerate viewing a fixation cross for 6.5 min, but six rs-fMRI runs were acquired instead of four, for a total of 21 min [47].

The HCP minimal preprocessing pipeline [48] was used. Briefly, several covariates were regressed from the data including linear and quadratic drift, mean cerebral-spinal fluid signal, mean white matter signal, and overall global signal. Images are bandpass filtered between 0.008 Hz and 0.12 Hz. As head motion can potentially confound fMRI, the mean framewise displacement [49] was calculated for each run for each participant and runs with a mean framewise displacement greater than 0.3 mm was removed from further analysis. Additionally, iterative smoothing, regression of 24 motion parameters (6 rigid-body parameters, 6 temporal derivatives of these parameters, and these 12 parameters squared) and frame censoring (displacement>0.3 mm) was performed to minimize motion confounds.

For each participant, we concatenated their rs-fMRI data from the first day for analysis and the data from the second day for validation. For each-day dataset, we excluded the participants’ runs whose rs-fMRI contained >30% frames flagged as motion outlier. Therefore, 528 participants (age range in month: 67-263) were left for the analysis. Among them 290 were females; and 521 participants had second day data for validation, (286 females). Detailed information including participants’ age distribution by sex and number of frames can be found in the Supplementary file (Figures S1, S2). Based on the Cole et al. parcellation [50] which contains 718 cortical and subcortical regions of interest (ROIs), the time course of each ROI in each participant was computed as the average time course across all voxels in the ROI. Further, we created ROI-to-ROI correlation matrices ***Σ***_***p***×***p***_, i.e., the FC matrices, using standardized fisher z-transformed Pearson correlation between timeseries.

### 4.3. Identification of age-specific patterns of FC development

As FC networks do not necessarily follow a linear association with age, we propose a sliding-window based clustering approach to identify a refined age interval of development for the brain regions and function networks. Suppose there are *N* participants with resting-state time sequence of *p* ROIs and their FC matrices 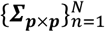. The (*i, j*)th entry of a FC matrix ***Σ***_*ij*_ represent the functional connections between ROI *i* and *j*. Since ***Σ***_***p***×***p***_ is symmetric, the following analyses are applied on the upper triangular entries of ***Σ***. As illustrated in Fig 1., our approach consists of two main steps: first, use sliding-window regression to capture age-related changes for each pair of the functional connections; second, use clustering method to identify clusters of functional connections that have similar changing curves. For each cluster, the cluster center characterizes the one type of FC and age association curves, and the corresponding indices matrix of the functional connections indicates the FC pattern of the cluster.

Step 1. We use sliding-window regressions to delineate the dynamic FC-age relationships. As shown in Figure S1, the number of participants in [67,100] is too small and will make the estimation unstable, thus we set fix age points in months from *T*_*begin*_ = 100 to *T*_*end*_ = 260 with interval *T*_*int*_ = 5, i.e., {*T* | *T* ∈ [100, 260] ∧ *T* = 5*x, x* ∈ *N*^+^}. To be more specifically, for a given window [*t* − *w, t* + *w*], *w* is the half window size and 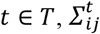 denotes the functional connections between ROI *i* and *j* that falls in the given window, we fit the following linear regression model

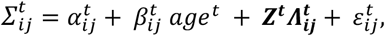

where the response variable 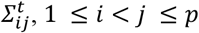 is the (i, j)th entry of the upper-triangular FC matrix in the given window, *age*^*t*^ ∈ [*t* − *w, t* + *w*], ***Z***^***t***^ is covariates in the given window. The regression coefficient sequence 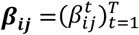 quantifies the changing direction and intensity at different age point. The regression coefficient sequence map is denoted as ***B*** = {***β***_*ij*_}_1≤*i*<*j*≤*p*_

Step 2. We apply the K-means clustering [51] on ***B*** into **K** sets ***S*** = {*S*_1_, *S*_2_, …, *S*_*K*_} to identify similar developing trends by minimizing the within-cluster sum of squares

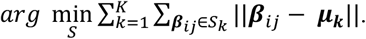

Each cluster center ***μ***_***k***_ = (*μ*_*k*1_, *μ*_*k*2_, …, *μ*_*kT*_), where *k* = 1,2, …, *K*, representing one type of FC-age changing curve, is then estimated through b-spline regression:

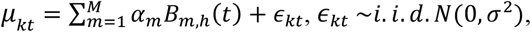

where *h* is the order of the spline function and *M* is the number of spline basis functions.

For each cluster *S*_*k*_, *k* = 1,2, …, *K*, we define its corresponding FC pattern 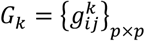, where 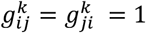, if 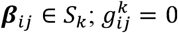, otherwise.

#### Parameter selection

The order of the spline function is set to be 4, which gives cubic splines. To determine the best window size *w*, we use the mean adjusted R-square [52] to measure the goodness of fit of sliding window regressions and select the optimal *w* through the elbow rule. The optimal number of spline bases *M* is selected by minimizing the mean of generalized cross-validation (GCV) criterion [53] that is defined as

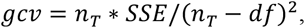

where *n*_*T*_ is the number of observations in the sequence, which in our case *n*_*T*_ = 33, *df* is the degrees of freedom measure of the smooth, and SSE is the error sums of squares. For the K-means clustering, the optimal number of *K* is selected through the elbow rule and the center of each cluster is acquired through 100 iterations.

The basic assumption in this paper is that FC changes nonlinearly with age. We employ the rolling regression/correlation techniques [54, 55] to delineate the curves with dynamic parameters, which is a local linear regression model. Compared with other curve fitting methods, it has the following advantages. First, unlike some nonlinear regression models (e.g., b-spline regression), the interpretation of the parameters is straightforward. Second, the estimation of the curve values is not the main interest, but the gradients that represent the changing speed. The local linear regression model has a direct estimation.

### 4.4. Sliding-window regression modelling

#### Delineate age-related changes in FC separately by sex

Previous studies shows that male and female have pronounced difference in development [56, 57]. To this end, we fit the model separately to acquire their age-related curves. For each sex *G* and a given window [*t* − *w, t* + *w*],

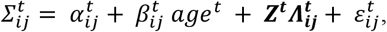

where ***Z*** represents the covariates including race (White, Asian, Black or African American), ethnicity (Hispanic or non-Hispanic), percentage of frames identified as motion outliers, mean framewise displacement, and site.

#### Compare sex differences in FC changing rate

We considered the effect of age and sex interaction on FC in the sliding-window regression analysis. For a given window [*t* − *w, t* + *w*],

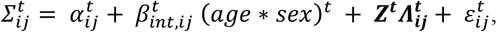

where ***Z*** represents the covariates including age, sex, race, ethnicity, percentage of bad frames, mean framewise displacement, and site information. The 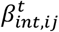 is the FC difference in slope of age between male and female. In our setting, a positive value of 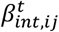 indicates that the slope in male is higher.

The optimal parameter set (w,M,K) for each case are (20,19,7) for female, (20,20,7) for male, and (20,11,4) for sex comparison.

### 4.5. Summary statistics of patterns of FC development

We summarize patterns of FC development from 3 levels: global measures, modular connectivity, and local hubs.

#### Global measures

We consider the network density, transitivity, and global efficiency to describe the FC patterns [58]. The transitivity measures the ratio between the number of triangles and the number of connected node triples that indicates the brain functional segregation, i.e., the ability for specialized processing to occur within densely interconnected groups of brain regions. The global efficiency is the average inverse shortest path length that indicates the ability to combine specialized information from distributed brain regions, also known as the functional integration ability.

#### Modular connectivity

The 718 brain regions can be divided into 12 functional networks (FNs) to study regional connectivity, including visual 1 network (V1N), visual 2 network (V2N), somatomotor network (SMN), cingulo-opercular network (CON), dorsal attention network (DAN), language network (LAN), fronto-parietal network (FPN), auditory network (AUD), default mode network (DMN), posterior-multimodal network (PMN), ventral-multimodal network (VMN) and orbito-affective network (OAN). We further separated the nodes by the anatomical location on cortical or subcortical surfaces and get 24 FNs (12 on cortical surface and 12 subcortical). We count the frequency of the connectivity’s in each intra-/inter-modules and conduct hypergeometric tests at significance level *α* = 0.05 with FDR correction.

#### Hubs

We analyzed hub nodes to gain more insights into the important ROIs of each development pattern. Here we define hubs as nodes (ROIs) with degrees at least three standard deviations higher than the mean degrees [59].

## Data availability

The human connectome project-development (HCP-D) data are publicly available [22]. (https://www.humanconnectome.org/study/hcp-lifespan-development)

## Code availability

The proposed method was implemented with R and the code is available upon request.

## Acknowledgements

This study was supported by the National Institute of Health (R01MH124106).

## Author Contributions

**Aiying Zhang:** Conceptualization, Formal analysis, Investigation, Methodology, Software, Validation, Visualization, Writing - original draft, Writing - review & editing; **Seonjoo Lee:** Conceptualization, Methodology, Project administration, Resources, Data curation, Supervision, Funding acquisition, Writing - review & editing; **David Pagliaccio:** Conceptualization, Interpretation, Writing - review & editing; **Rachel Marsh**: Supervision, Interpretation, Writing - review & editing

